# The Impact of Population Stratification on the Analysis of Multimodal Neuroimaging Derived Measures

**DOI:** 10.1101/2022.08.06.503037

**Authors:** Tzu Hsuan Huang, Robert Loughnan, Wesley K Thompson, Chun Chieh Fan

**Affiliations:** Center for Human Development, University of California, San Diego; Laureate Institute for Brain Research, Tulsa, Oklahoma; Division of Biostatistics and Bioinformatics, School of Public Health, University of California, San Diego; Department of Radiology, School of Medicine, UCSD

**Keywords:** Magnetic resonance imaging, Adolescent Brain Cognitive Development Study, Brain imaging features, Population stratification, Genetic ancestry factors, imaging confounds, Parental income, task fMRI, rsMRI, DTI, RSI, sMRI

## Abstract

Magnetic resonance imaging (MRI) studies of the human brain are now attaining larger sample sizes with more diverse samples. However, population stratification, a key factor driving heterogeneity and confounding of associations, is seldom accounted for in neuroimaging analyses. To investigate this issue, we assessed the impact of population stratification on multimodal imaging measures using baseline data from the Adolescent Brain Cognitive Development (ABCD) Study^SM^ (n = 10,748). Given this sociodemographically diverse sample, which broadly reflects the population composition of the United States, we performed a thorough evaluation of the impact of population stratification on derived neuroimaging metrics across five different imaging modalities: task functional MRI (task fMRI), resting state functional MRI (rsMRI), diffusion tensor images (DTI), restricted spectrum images (RSI), and structural T1 MRI (sMRI). We used parental income level as an example to highlight the impact of population stratification in confounding brain-wide associations. We show that derived metrics from structural images have up to three times more signal related to population stratification than do functional images. Controlling for population stratification in statistical models leads to a substantial reduction in the association strength between variables of interests and imaging measures, indicating the scale of potential bias. Moreover, because of unequal access to resources (such as income) across ancestral groups in United States, population stratification effects on imaging features may bias associations between parental income levels and imaging features, as we demonstrate. Our results provide a guide for researchers to critically examine the impact of population stratification and to assist in avoiding spurious brain-behavior associations.

**Highlights:**

- Here, we conduct a comprehensive survey of the confounding impact of population stratification in large-scale imaging studies.
- Morphological features from structural imaging appear to be more susceptible to the confounding effects of population stratification than do functional imaging features.
- The population stratification tends to inflates the association strengths between the variable of interest and imaging features.
- When the variable of interest is highly colinear with the population stratification, such as income levels, brain associations cannot be differentiated and may be misattributed as mediating effects.
- It is critical to account for population stratification in imaging analyses.

## 1. Introduction

Magnetic resonance imaging (MRI) has become an essential tool for measuring neuroanatomical and functional variation of the human brain. Structural and functional imaging derived features have been used for understanding brain-behavioral associations (Bernanke J et al., 2022), neurodevelopment (Bethlehem RAI et al., 2022), and the purported impact of environmental exposures on the brain (Marshall AT et al., 2020). Although effect sizes and test-retest reliability of MRI features have recently been critically examined (Dick et al. 2021; Kennedy JT et al., 2022; Marek S et al., 2022), the internal validity of associations based on neuroimaging features has seldom been discussed. In particular, brain-behavior associations are susceptible to a well-known confound in genetic studies, population stratification, especially given that, as neuroimaging studies have become larger, they are beginning to include more diverse and heterogeneous samples.

Population stratification is defined as heterogeneity driven by sub-populations due to differences in genetic ancestral background (Tanaka H et al., 2021). Variation in genetic ancestry is largely explained by population history (Hartl & Clark, 2007) and can be correlated with cultural and social identities (Novembre et al., 2008). Ancestral features in the human genome often correlate with social and physical environments, despite the absence of any actual causal relationships (Price et al.,2006). To appropriately account for this, it is considered best practice to control for population stratification in genome-wide association studies (GWAS’s), reducing false positives driven by the potential correlation between genetic ancestry and social and physical environments (Dehghan A, 2018). However, to date high-profile brain-wide association studies still treat the potential confounding impact of population stratification as a secondary issue, if at all (Marek S et al., 2022).

Associations between genetic ancestry and structural imaging features have been reported in the literature. The shape of cortical folding, measured by structural T1 images, was shown to be associated with genetic ancestry across four major continental groups in a large-scale imaging cohort of the US, despite no evident associations with the functionally relevant measures among participants (Fan et al., 2015). Intra-cranial volume and total surface area have been shown to significantly associate with genetic clines within European cohorts (Bakken et al., 2011), indicating that structural imaging measures can be sensitive to even fine-level variation in the population substructure. However, there has been no in-depth investigation into the relationship between population stratification and multi-modal imaging measures. For example, it is currently unclear the degree to which population stratification associates with neuroimaging features beyond metrics derived from structural T1 images, such as microstructural integrity from diffusion-weight images, task-related variations in the BOLD signal, and resting state functional connectivity.

Furthermore, population stratification is often correlated with environmental disparities between individuals (e.g., due to structural racism), disparities which may be associated with unequal outcomes or which may have neurodevelopmental consequences on brain structure and function. Many studies have reported that socioeconomic disparities (e.g., parental income) influence brain development and cognitive functioning (Hackman & Farah, 2009; Taylor et al., 2020; Hackman et al., 2021). In particular, parental income was significantly associated with cortical thickness and total surface area (Noble et al., 2015). Poverty has also been linked to structural differences in several areas of the brain (e.g., cortical thickness, white and cortical gray matter volume, hippocampal size and amygdala volumes) and associated with learning and cognitive abilities among children (McLoyd, 1998; Luby et al., 2013; Lawson et al., 2013; Hair et al., 2015; Merz et al., 2020). Living in poor neighborhoods has been associated with high risk of exposure to environmental toxins in addition to differences in total brain volumes and cortical surface measures (Marshall et al., 2020).

Population stratification can bias imaging analyses in several ways. First, a given imaging feature may associate with population stratification due to cranial morphology without any functional consequences. If population stratification also correlates with the variable of interest, such as parental income, morphological imaging features would be found to be associated with the variable of interest despite no true effect existing between images and the outcome of interest (Figure 1a, pure confounding effect). Second, there may be an actual (non-zero) association between the variable of interest and the neuroimaging features, but the estimated relationship may be biased upwards or downwards due to conflation of true effect and population stratification (Figure 1b, inflated estimation driven by confounds). Third, population stratification and the outcome of interest may be so correlated that its effects cannot be differentiated in an observational study (Figure 1c, misattribute as mediations).

**Figure 1.**
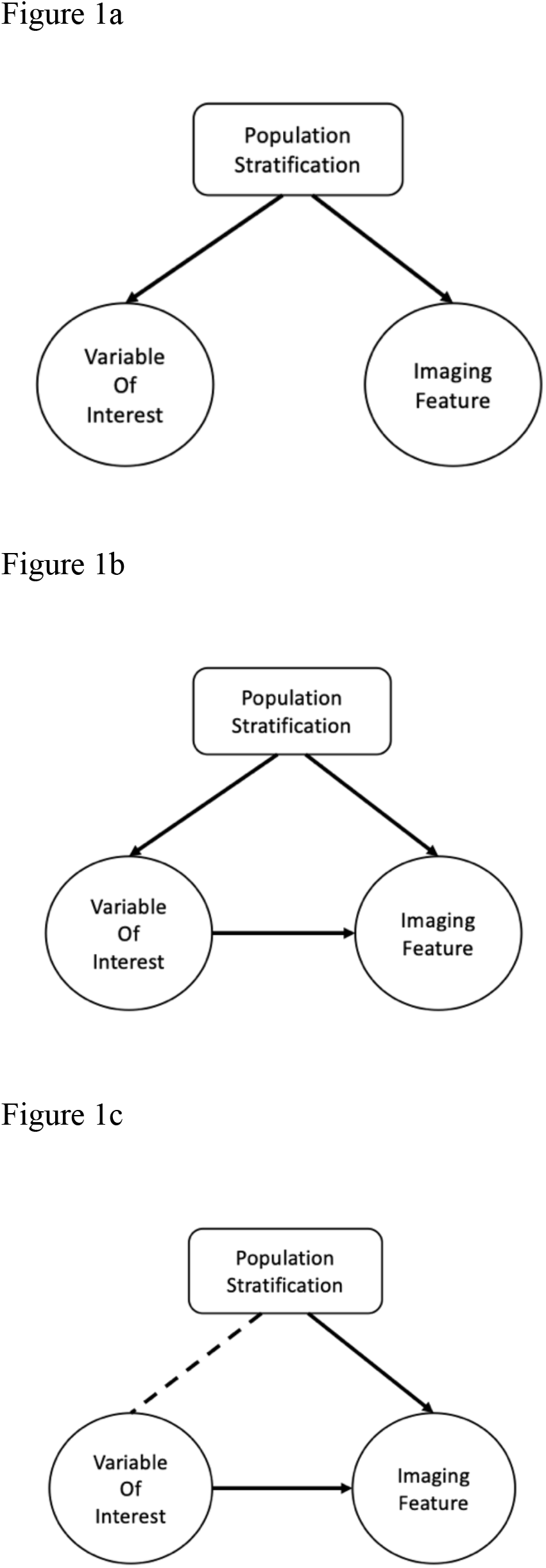
Three roles of the population structure between income and imaging measures. **Figure 1a. Pure confounding effect**. Assuming imaging feature is associated with population stratification and the variable of interest in the study is correlate to population stratification. Under the circumstances, the imaging feature can be found to be associated with income despite no actual causal relationships. **Figure 1b. Inflated estimation driven by confounds**. Assuming the variable of interest in the study has effect on imaging feature, imaging feature is associated with population stratification, and the variable of interest in the study is also correlate to population stratification. The estimation of the relationship between variable of interest and imaging feature is biased due to the conflation of true effect and population stratification. **Figure 1c. The misattributed mediations**. Assuming population stratification and the variable of interest can be so intertwined. The true effect of variable of interest and confounding effect of population stratification are not differentiable. The effect of population stratification would mask the true associations involved the variable of interest and over-attribute the source of variations to the population stratification.

As more imaging studies strive to enroll participants from diverse backgrounds and reach population-relevant scales, understanding the impact of population stratification on variability in neuroimaging metrics becomes increasingly important. To raise the awareness of this issue, we critically examined the aforementioned three possible confounding scenarios in imaging analyses using the large-scale, ancestrally, and socioeconomically diverse participants from the Adolescent Brain Cognitive Development^**SM**^ (ABCD) Study. Based on the ABCD Study® data, we show that population stratification has stronger associations with structural measures than with functional measures derived from MRI. Furthermore, we demonstrate that population stratification can significantly bias associations between parental income and brain imaging features, regardless of the imaging modality. The strong correlations between population stratification and income in this observational study makes many of the effects between ancestral features and structural metrics impossible to be differentiate from the true effects of the income differences. Using this example, we illustrate the importance of confounding effects due to population stratification and to caution researchers analyzing and interpreting results from brain-behavioral associations in diverse samples about this important source of confounding bias.

## 2. Methods

### 2.1. ABCD data

The ABCD Study® is a longitudinal study that recruited n=11,878 healthy children aged 9-10 years old at baseline. Participants have a high degree of socio-economic and demographic diversity, broadly reflecting the US population in this age group (Dick et al, 2021). The ABCD Study will follow their development of for ten years with yearly in-person assessments. Data are released publicly on an annual basis via the National Institute of Mental Health Data Archive (NDA, https://data-archive.nimh.nih.gov/abcd). In this study, we excluded individuals who have missing information due to ascertainment missingness (missing demographics info), imaging acquisition problems (missing imaging region-of-interest measurements), and genotyping data failures (cannot infer population stratifications). The resulting sample of 9,694 individuals was used in these analyses. Demographic characteristics can be found in the Table 1.

**Table 1.** Summary of ABCD baseline demographics in this study.

|  | N (%) | Mean (SD) |
| --- | --- | --- |
| **Age** | 9,694 | 9.92 (0.6) |
| **Sex** |  |  |
| Male | 5,058 (52.2) |  |
| Female | 4,636 (47.8) |  |
| **Parental Income** |  |  |
| [< 50K] | 2,737 (28.2) |  |
| [>=50K & <100K] | 2,790 (28.8) |  |
| [>= 100K] | 4,167 (43.0) |  |
| **GAF** |  |  |
| Pop1(%) |  | 17 (0.3) |
| Pop2(%) |  | 11 (0.2) |
| Pop3(%) |  | 4 (0.2) |
| Pop4(%) |  | 68 (0.4) |

GAF: Genetic Ancestry Factors; Pop1: African Ancestry; Pop2: Native American; Ancestry Pop3: East Asian Ancestry; Pop4: European Ancestry. Summarize genetic variation consistent with the four ancestral components, representing the degree of genetic similarity between individuals and the four continental populations. Sum of the proportion of GAF across four continental groups is one.

### 2.2. Measures

#### 2.2.1 Metrics for population stratification

ABCD participants were genotyped on using the Affymetrix smokescreen array (Baurley et al., 2016). After basic quality controls (success rate of genotype calls and overall missingness rate under 10%), 516,598 genetic variants were placed into fastStructure to infer population stratification (Raj et al., 2014). Based on information criteria, the best fitting model for the ABCD sample indicates four major population groups: African ancestry, Native American ancestry, East Asian ancestry, and European ancestry. Proportion of genetic ancestry for each population groups was coded in genetic ancestry factors (GAFs) for each participant, with values between 0 and 1. As these factors typically represent confounds due to gene by environment correlation, we label these components as Pop1 (African), Pop2 (Native American), Pop3 (East Asian), and Pop 4 (European) in analyses to emphasize they are not meant to be interpreted out of context. Because the proportions across all four GAFs sum to unity and Pop4 has the largest number of average proportions, we dropped the Pop4 in all of our analytic models, treating it as the reference category.

#### 2.2.2. Multimodal Imaging Measures

The multimodal imaging features used in this report are derived region-of-interest (ROI) measures from NDA data release 3.0. Neuroimaging data were harmonized across 21 sites and processed by the ABCD Data Analysis Informatics and Resource Center (DAIRC) and the ABCD Image Acquisition Workgroup (Hagler et al., 2019). Derived ROI measures were obtained from the five MRI modalities available in the ABCD data release: 1) structural T1 MRI (sMRI, including cortical surface derived measures in Desikan atlas as surface area, surface thickness, sulcal depth, and subcortical volumes); 2) diffusion tensor images (DTI, including fractional anisotropy and mean diffusivity across major fiber tracts); 3) restricted spectrum images (RSI, normalized restricted isotropic and restricted anisotropic across both cortices, subcortical regions, and major fiber tracts). 4) task functional MRI (task fMRI, average BOLD signal consists of contrasts from the N-Back task, Monetary Incentive Decision task, and Stop-Signal Response task in Desikan Atlas); and 5) resting state functional MRI (rsMRI, derived connectivity measures across Gordon’s parcellation and subcortical regions, summarized in network partitions (Gordon et al., 2017)). Based on these selection criteria, there were a total 6,641 multimodal measures used in this study, where the number of features for each modality were: 1) 1,184 sMRI; 2) 2,376 DTI; 3) 1,754 RSI; 4) 891 task fMRI; and 5) 436 rsMRI. Details of the imaging acquisition, processing pipelines, and quality control metrics leading to the release of the ABCD imaging dataset on NDA can be found in prior publications (Hagler et al., 2019; Casey BJ et al., 2018).

### 2.3. Statistical Analysis

We examined three possible confounding scenarios, as illustrated in Figure 1. In the first scenario, we quantified the maximum amount of confounding population stratification could give arise to, assuming there were no true effects of the variable of interest on the imaging measures (Figure 1a). Here, we performed Likelihood Ratio Tests (LRTs) to investigate whether GAFs were associated with imaging measures. Models included the fixed effects of sex at birth, age, scanner serial numbers, and GAFs (Pop1-Pop3), compared to a base model without including GAFs. All models controlled for family membership via random effects. Variance explained was computed based on the differences in the pseudo-R^2^ from the LRT of the full model vs. the base model.

In the second scenario, we assessed how much effects could be inflated by population stratification, using parental income at baseline as an example (Figure 1b). The total parental income reported by the guardians during the first assessment. We again fit two linear mixed models, one with and the other without controlling GAFs. Both models included fixed effects of sex at birth, age, parental income, and scanner serial number. We assessed the confounding impact of GAFs on the association between parental income and brain measures by comparing the proportion of “significant” (p < .05) parental income regression coefficients of these two models and by the percentage changes of the coefficients after inclusion of GAFs. All analyses again controlled for family membership via random effects.

In the third scenario, we assessed how much associations of GAFs with brain measures carry over to parental income demonstrating the difficulty in differentiating effects of population stratification from those of the variable of interest (Figure 1c). To achieve this goal, we use mediation analysis (Imai K et al., 2010) to compute the proportion of the effect from GAFs to imaging measures explained by the covariation of GAFs with parental income levels. Note, while we use the methods of mediation, we are not positing a causal model; rather, we are estimating “carry over” effects, computed using these same methods. Thus, we use parental income as an example and interpret the effects computed from the mediating model into 1) the effect of GAFs on ROIs after partialling out the effect of parental income and also 2) the effect of GAFs on ROIs carries through by parental income. As before, all mediation models control for age, sex, the serial number of the imaging device, and family membership.

To evaluate the likelihood that we would find significant associations for a given imaging modality, we first calculated the proportion of ROIs reaching a nominal significance threshold (p-value < 0.05) of effect of GAFs on ROIs, effect of GAFs on ROIs after partialling out the effect of parental income, and the effect of GAFs on ROIs carries through by parental income. In order to compare differences between each imaging modality, we used task fMRI as the reference category, since it consistently has the lowest proportion of ROIs reaching the nominal significance threshold. We then computed the risk ratio (RR) for significance for each modality, i.e., the ratio of the probability of reaching nominal significance threshold given the imaging modality, representing how much more or less likely that modality is to pass the nominal 0.05 significance threshold than the reference imaging modality. We choose the task fMRI as the reference. RR allows for intuitive comparison of the sensitivity of each imaging modality for different ancestral backgrounds. We estimated 95% confidence intervals of RR via 10,000 bootstrapping iterations, repeatedly sampling participants with replacement. Mixed models and mediation analysis were implemented using the R *lme4* and *mediation* packages (Tingley et al., 2014).

## 3. Results

### 3.1 Pure Confounding Effect

Table 2 displays the percentage of significant associations and the distribution of the variance explained (R^2^) of GAFs with ROIs across the five imaging modalities; sMRI had the highest significance rate (87.8%) with GAFs, follow by structural connectivity measures (DTI=80.9% and RSI=69.1%). Functional measures had lowest significance rate (rsMRI =59.3% and task fMRI =25.2%). The distribution of the variance explained is consistent with the significance rates, as the sMRI has a higher variance explained than other imaging modalities.

**Table 2.** Pseudo-R2 of Likelihood Ratio Test to investigate mixed model fit improvement with GAFs in the model, among the significant ROIs. *Based on Nagelkerke’s pseudo-R2

|  | Percentage of<br>significant in<br>LRT | Distribution of variance explained* |  |  |  |
| --- | --- | --- | --- | --- | --- |
|  |  | Q1 | Median | Q3 | Max |
| **task fMRI** | 25.2% | 0% | 0.1% | 0.2% | 3.3% |
| **rsMRI** | 59.3% | 0.1% | 0.1% | 0.2% | 3.0% |
| **RSI** | 69.1% | 1.7% | 2.7% | 3.6% | 6.3% |
| **DTI** | 80.9% | 1.7% | 2.7% | 3.7% | 6.3% |
| **sMRI** | 87.8% | 1.7% | 2.5% | 4.4% | 13.2% |

### 3.2 Inflated Estimation Driven by Confounds

In ABCD cross-sectional samples, GAFs were significantly associated with parental income (Pop1: β=-2.58, p-value <0.000; Pop2: β=-3.01, p-value <0.000; Pop3: β=0.07, p-value=0.838) with a Nagelkerke Pseudo-R^2^ of 0.332. Given the strong associations between population stratification and parental income, we used parental income as an example of how much population stratification can bias imaging analyses. Table 3 shows the percentage of significant tests of parental income regression before and after controlling GAFs, and the percentage coefficient change after controlling for GAFs (difference in association coefficients with the coefficient before controlling for GAFs as the denominator). The percentage of significant associations of parental income with ROIs were markedly lower after controlling for GAFs for all imaging modalities (Table 3), but especially for sMRI (median percentage coefficient change of 52%), followed by structural connectivity measures (DTI=41.3%; RSI=33.6%), and rsMRI (38.0%). Task fMRI had the lowest coefficient changes (18.8%).

**Table 3.**
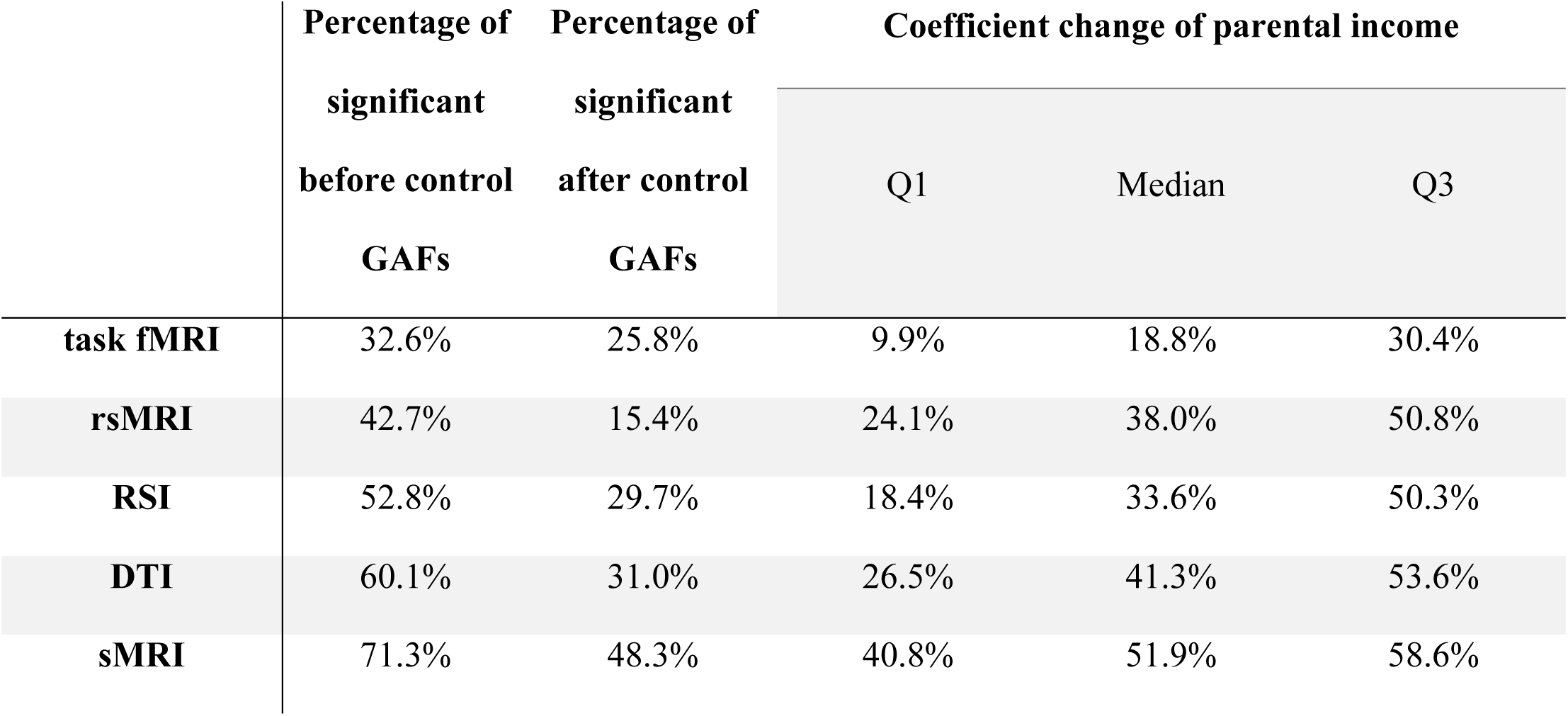
Parental income regression coefficients percentage change from the regression model change due to GAFs, among the significant ROIs. GAF: Genetic Ancestry Factors.

### 3.3 The misattributed mediations

In the third scenarios, we decomposed the correlated effects of parental income and GAFs on ROIs. The left column of Table 4 shows the significance proportions of misattributed mediation effects of GAFs on ROIs by parental income, using average causal mediation estimation model, and also displays the proportion of variance misattributed among the significant ROIs. sMRI had highest significance rate (50.8%) of ROIs carryover by parental income follow by DTI (35.0%), RSI, rsMRI (34.4%) and task fMRI (27.7%). Among ROIs which has significant associations with parental income, RSI (median =24.8%) has the largest proportion of carryover effects in general and follow by sMRI (median =23.3%), DTI (median=21.7%), and rsMRI (median =21.8%). Task fMRI, on the other hand, has the least (median=13.2%). These results reflect the volume of the carryover effect when the effect of GAFs travel with parental income. The significance rates across imaging modalities in the analyses 1 to 3 are summarized in the Figure 2.

**Table 4.**
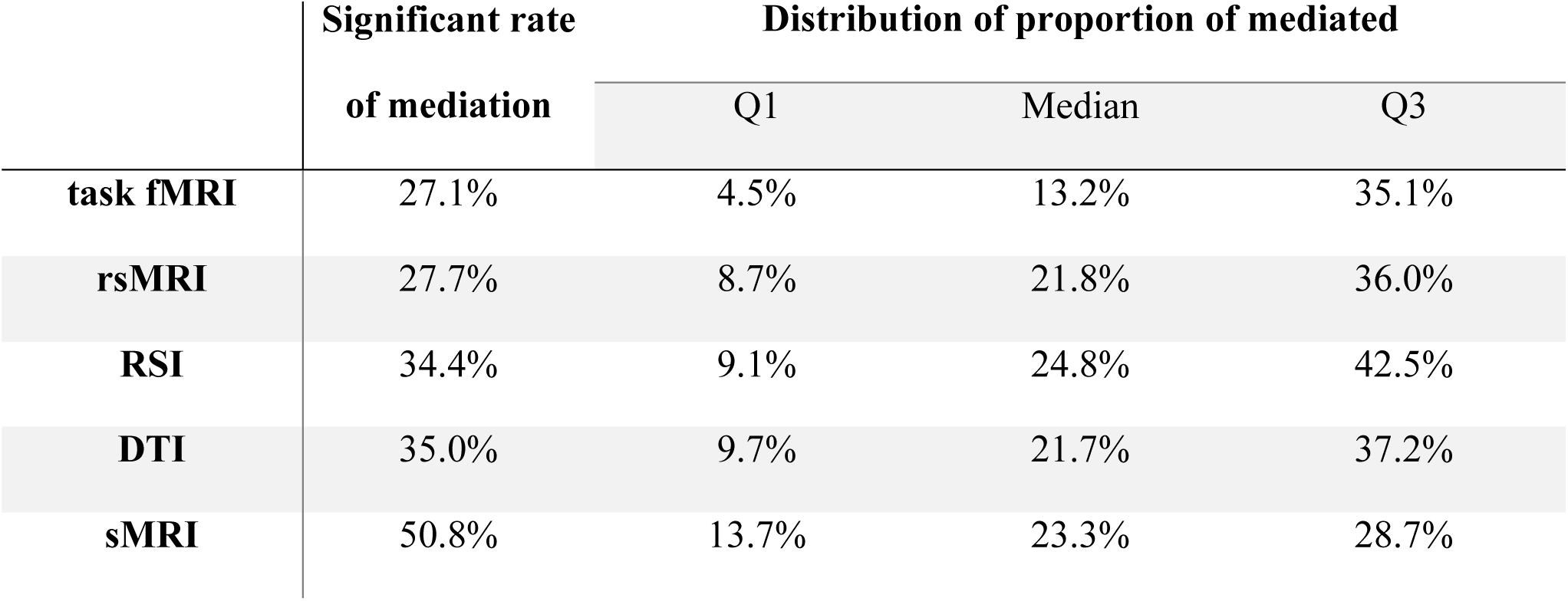
Description of proportion of mediated in association between neuroimaging and GAFs through Parental income for each ROIs, among the significant ROIs.

|  | Significant rate | Distribution of proportion of mediated |  |  |
| --- | --- | --- | --- | --- |
|  | of mediation | Q1 | Median | Q3 |
| **task fMRI** | 27.1% | 4.5% | 13.2% | 35.1% |
| **rsMRI** | 27.7% | 8.7% | 21.8% | 36.0% |
| **RSI** | 34.4% | 9.1% | 24.8% | 42.5% |
| **DTI** | 35.0% | 9.7% | 21.7% | 37.2% |
| **sMRI** | 50.8% | 13.7% | 23.3% | 28.7% |

**Figure 2.**
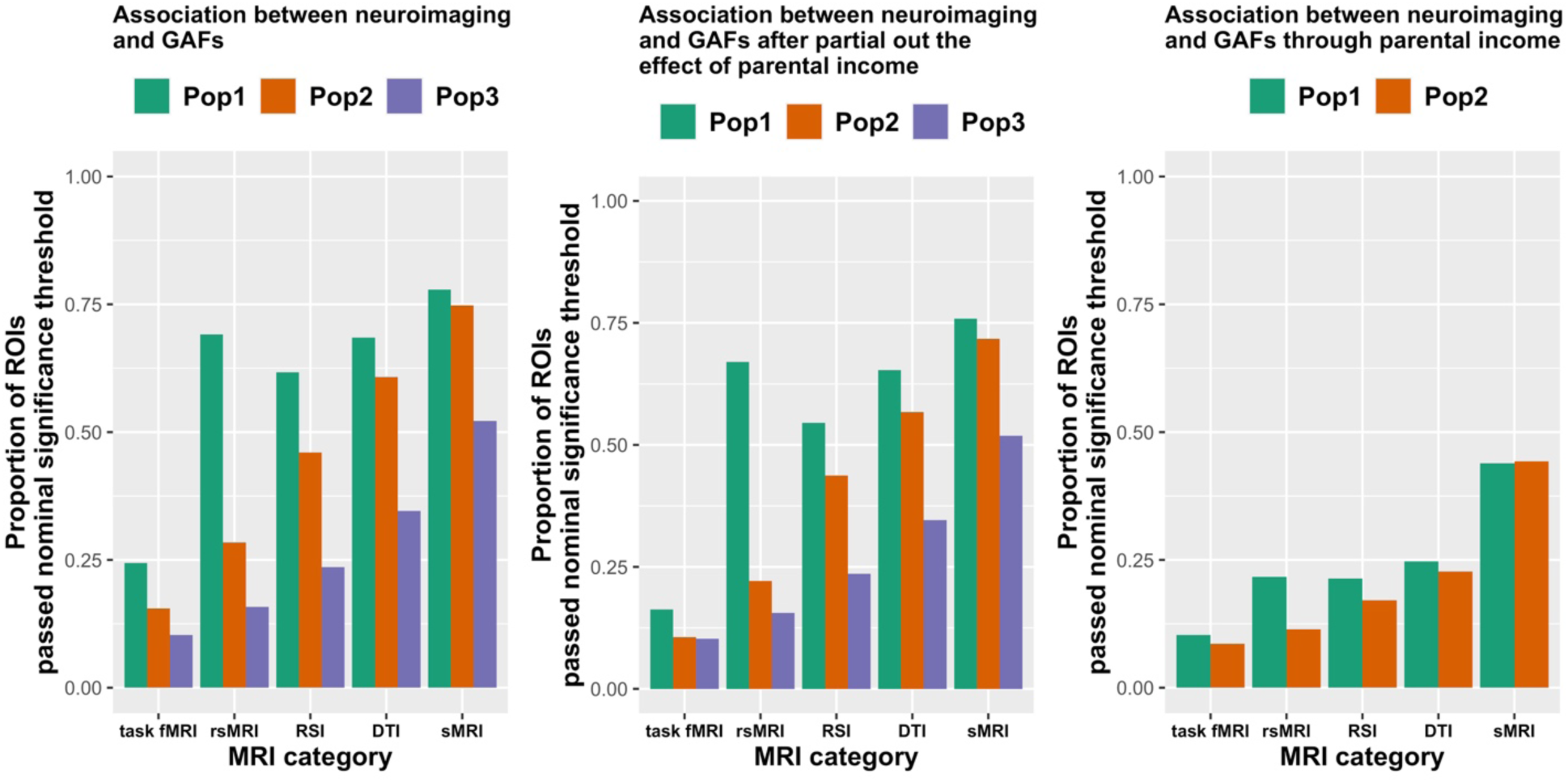
Proportion of ROIs passed nominal significance threshold in mediation analysis, stratified by MRI category and genetic ancestry factors. These three panels show the proportion of ROIs passed nominal significance threshold in association between neuroimaging and GAFs, association between neuroimaging and GAFs after partialing out the effect of parental income and association between neuroimaging and GAFs through parental income (from left to right). The color of the bar represented the genetic ancestry; Green: Pop1; Orange: Pop2; Purple: Pop3. * Pop4 was not estimable in the average causal mediation effect due to small sample size limitation.

### 3.4. The risk of finding spurious associations

To more formally assess the qualitative trends for each modality in the proportion of ROIs reaching a nominal significance threshold and for a more intuitive observation, we compared the other four imaging modalities against task fMRI to assess the proportion of ROIs reaching a nominal significance threshold by calculating their risk ratios (RR) of finding associations. In Figure 3, all four imaging modalities (sMRI, DTI, RSI, rsMRI) show the increased likelihood (RR > 1) of finding associations compared to task fMRI. The ranking (in order of likelihood of finding significance) was 1) sMRI; 2) DTI; 3) RSI; 4) rsMRI; and 5) task fMRI. After partialling out parental income, the association with GAFs (orange) for Pop1 and Pop2 were all stronger than the total association with GAFs (green). Additionally, sMRI had the strongest association both with i) GAFs after partialling out the parental income (orange, Pop1 = 4.7; Pop2 = 6.9; Pop3 = 5.1, Figure 3, Supplemental Table 1) and ii) GAFs carried through parental income (purple, Pop1 = 4.3; Pop2 = 5.2, Figure 3, Supplemental Table 2), having at least four times likely to be claimed as significant comparing to task fMRI. Moreover, sMRI is the only imaging modality with stronger association through parental income than overall association (green, Pop1 = 3.2; Pop2 = 4.9, Figure 3, Supplemental Table 1).

**Figure 3.**
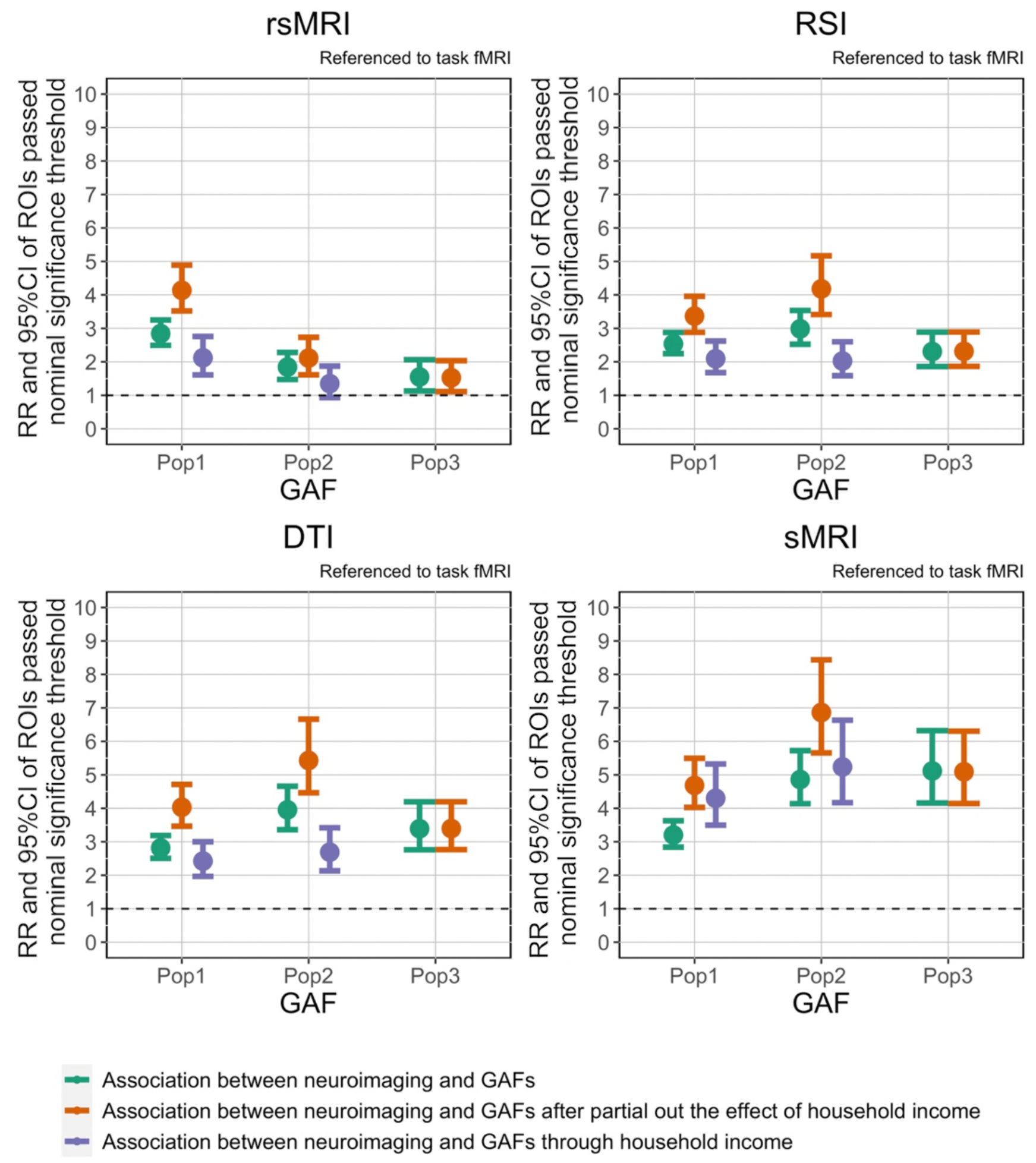
The RR and 95%CI of ROIs passed nominal significance threshold in mediation analysis (referenced to task fMRI). All four imaging modalities (sMRI, DTI, RSI, and rsMRI) were referenced to task fMRI and calculating relative risk (RR) and 95%CI of ROIs passed nominal significance threshold in mediation analysis stratified by genetic ancestry factors. The mean of RR and its 95% confidence interval were shown in the bar plots. The green bar represented the mean and the 95% confidence interval of RR of association between neuroimaging and GAFs; The purple bar represented the mean and the 95% confidence interval of RR of association between neuroimaging and GAFs through parental income; the orange bar represented the mean and the 95% confidence interval of RR of association between neuroimaging and GAFs after partial out the effect of parental income. * Pop4 was not estimable in the average causal mediation effect due to small sample size limitation. *GAF: Genetic Ancestry Factors; Pop1: African Ancestry; Pop2: Native American; Ancestry Pop3: East Asian Ancestry; Pop4: European Ancestry.

## 5. Discussion

Our results show that structural T1 imaging measurements exhibited the strongest associations with population stratification in terms of magnitude and probability of significant associations in the ABCD Study sample. Functional brain measures, especially task fMRI, showed limited or negligible associations with population stratification. This reaffirms previous conjectures that the impact of population stratification on imaging measures is attributable to differences in the morphological features with little to no functional relevance (Fan et al., 2015). Analyses on the connectivity measures show the same trend, as the functional connectivity have much weaker association signals than the structural connectivity (DTI and RSI).

Recent reports on shared genetic loci between structural imaging measures and cranial features have indicated shared molecular origins, driven by the closely coordinated developmental processes of ectoderm and neural ectoderm (Naqvi et al., 2021). This suggests that structural measures from neuroimaging may be particularly susceptible to the confounding effects of population stratification. Given that over 90% of structural ROIs are associated with GAFs and highly likely to lead to misattribution of the impact of variable of interests, any analysis using structural images should seriously consider the biasing impact of population stratification.

Although most evident in sMRI associations, the confounding and carryover effects of population stratification are ubiquitous across imaging modalities. As illustrated in our example analyses with the parental income using ABCD data, median changes of regression coefficients between parental income and imaging ROIs before and after controlling for population stratification are 18.8%, 38.0%, 33.6%, 41.3%, and 51.9% for imaging features from task fMRI, rsfMRI, RSI, DTI, and structural T1 MRI, respectively. The median proportion of significant carryover effects range between 13.2% to 23.3%, indicating non-negligible conflation between population stratification and parental income levels. This makes the interpretation on the brain-behavioral associations challenging, as controlling population stratification can take away some proportion of the true associations of the parental income, yet the correlation of parental income with population stratification makes it impossible to differentiate true effects and confounds.

Our analyses do not comprehensively demonstrate the potential causes underlying the association of population stratification and neuroimaging metrics. Given the broad interest on the social determinants of variability in neuroimaging metrics (Noble et al., 2015; Gonzalez et al., 2020), there is a need to conduct bias analyses on the impact of population stratification on brain-behavior associations of interest to determine to extent to which this may confound effects detected. Researchers should be thoughtful when considering which covariates to include in any given analyses and be able to motivate the choices with theoretical and meaningful rationales.

This study has several limitations. First, we analyzed ROI-level data and contrasted between imaging modalities. We did not emphasize whether specific ROIs were more susceptible to population stratification across different modalities and dig into finer resolutions beyond these anatomically defined regions. Moreover, it is possible to construct a multivariate model that is highly predictive of population stratification despite weak associations at the individual ROI level, which emphasizes that population stratification may have small but distributed effects across the brain (Altmann & Mourao-Miranda, 2019). In this paper we focused on providing an overview of the associations rather than providing a general modeling strategy to account for confounding relationships between brain features and population stratification. Second, for simplicity we used a very rough definition of parental income (i.e., total parental income reported by the guardians during the first assessment) in our analyses.

There are several components of socioeconomic status, such as parental education, parent occupation and income-to-needs. Parental income does not fully represent the effect of socioeconomic status on the development of children’s brain structure. Third, we used GAFs to represent the population stratification, yet there are other measures of population structure, such as using genetic principal components (Patterson et al., 2006). Nevertheless, controlling for GAFs or for the first several genetic PCs typically give consistent results. We chose GAFs for the ease of presentation. Finally, we did not perform a full set of bias analyses to examine when and how population stratification can become a confounder or collider. While we believe formal bias analyses are critical, bias analyses are scenario/context specific, depending on which variables are used and which hypotheses are being tested.

Taken together, our results reveal the necessity for taking population stratification seriously in neuroimaging research. Given associations between population structure, self-declared race and ethnicity, and health disparities, it is imperative that these potential relationships are thoughtfully considered by researchers in the context of their analyses to avoid spurious brain-behavior associations especially where these may be stigmatizing for minority groups. These findings provide a valuable reference for future neuroimaging analyses using large-scale, diverse samples.

## Supporting information

Supplemantal Tabel1, Supplemantal Figure1

## Acknowledgements

This work was supported by grant R01MH122688 and RF1MH120025 funded by the National Institute for Mental Health (NIMH). Data used in the preparation of this article were obtained from the Adolescent Brain Cognitive Development SM(ABCD) Study (https://abcdstudy.org), held in the NIMH Data Archive (NDA). The ABCD Study® is supported by the National Institutes of Health and additional federal partners under award numbers U01DA041048, U01DA050989, U01DA051016, U01DA041022, U01DA051018, U01DA051037, U01DA050987, U01DA041174, U01DA041106, U01DA041117, U01DA041028, U01DA041134, U01DA050988, U01DA051039, U01DA041156, U01DA041025, U01DA041120, U01DA051038, U01DA041148, U01DA041093, U01DA041089, U24DA041123, U24DA041147. A full list of supporters is available at https://abcdstudy.org/federal-partners.html. A listing of participating sites and a complete listing of the study investigators can be found at https://abcdstudy.org/consortium_members/. ABCD consortium investigators designed and implemented the study and/or provided data but did not necessarily participate in the analysis or writing of this report. This manuscript reflects the views of the authors and may not reflect the opinions or views of the NIH or ABCD consortium investigators. The ABCD data repository grows and changes over time.

## Data Availability

The ABCD data used in this report came from https://doi.org/10.15154/1524729. The fast track data release used in this report are available at https://nda.nih.gov/edit_collection.html?id=2573. Instructions on how to create an NDA study are available at https://nda.nih.gov/training/modules/study.html).

