## Supplementary material for "The Impact of Population Stratification on the Analysis of Multimodal Neuroimaging Derived Measures": Supplemantal Tabel1, Supplemantal Figure1

### Table legends

**Table S1. RR and 95%CI of ROIs passed nominal significance threshold.**

### Figure legends

**Figure S1. MRI data flow chart.** Available MRI events (green) were divided into five modalities (blue) and trimmed incomplete or very poor-quality data (yellow). We further filtered to exclude the relative irrelevant ROIs (orange).

### Supplemental Materials

Table S1. RR and 95%CI of ROIs passed nominal significance threshold.

| Category | Genetic Ancestry Factors | $RR_{Total\ Effect}$[a] | 95%CI | $RR_{Direct\ Effect}$[b] | 95%CI | $RR_{Mediation\ Efffect}$[c] | 95%CI |
|---|---|---|---|---|---|---|---|
| fMRI | Pop1 | 1 | (1, 1) | 1 | (1, 1) | 1 | (1, 1) |
| fMRI | Pop2 | 1 | (1, 1) | 1 | (1, 1) | 1 | (1, 1) |
| fMRI | Pop3 | 1 | (1, 1) | 1 | (1, 1) | - | - |
| rsMRI | Pop1 | 2.8 | (2.5, 3.3) | 4.1 | (3.5, 4.9) | 2.1 | (1.6, 2.8) |
| rsMRI | Pop2 | 1.8 | (1.5, 2.3) | 2.1 | (1.6, 2.7) | 1.3 | (0.9, 1.9) |
| rsMRI | Pop3 | 1.5 | (1.1, 2.1) | 1.5 | (1.1, 2.0) | - | - |
| RSI | Pop1 | 2.5 | (2.3, 2.9) | 3.4 | (2.9, 4.0) | 2.1 | (1.7, 2.6) |
| RSI | Pop2 | 3.0 | (2.5, 3.5) | 4.2 | (3.4, 5.2) | 2.0 | (1.6, 2.6) |
| RSI | Pop3 | 2.3 | (1.9, 2.9) | 2.3 | (1.9, 2.9) | - | - |

| | | | | | | | |
|---|---|---|---|---|---|---|---|
| DTI | Pop1 | 2.8 | (2.5, 3.2) | 4.0 | (3.5, 4.7) | 2.4 | (2.0, 3.0) |
| DTI | Pop2 | 4.0 | (3.4, 4.7) | 5.4 | (4.5, 6.7) | 2.7 | (2.1, 3.4) |
| DTI | Pop3 | 3.4 | (2.8, 4.2) | 3.4 | (2.8, 4.2) | - | - |
| sMRI | Pop1 | 3.2 | (2.8, 3.6) | 4.7 | (4.0, 5.5) | 4.3 | (3.5, 5.3) |
| sMRI | Pop2 | 4.9 | (4.1, 5.7) | 6.9 | (5.7, 8.4) | 5.2 | (4.1, 6.6) |
| sMRI | Pop3 | 5.1 | (4.2, 6.3) | 5.1 | (4.1, 6.3) | - | - |

GAF: Genetic Ancestry Factors; Pop1: African Ancestry; Pop2: Native American; Ancestry Pop3: East Asian Ancestry; Pop4: European Ancestry;

a: The effect of GAFs on ROIs;

b: The effect of GAFs on ROIs after partial out the effect of parental income;

c: The effect of GAFs on ROIs carries through by parental income.

* Pop4 was not estimable in the average causal mediation effect due to small sample size limitation.

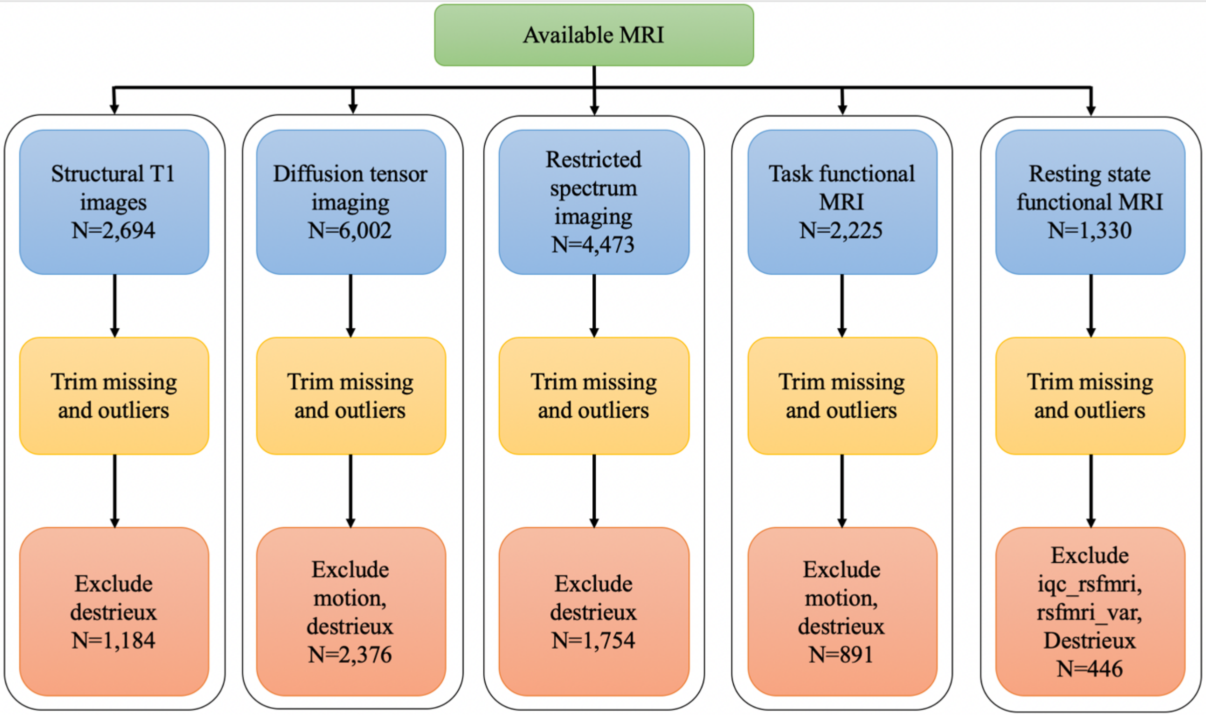


Figure S1. MRI data flow chart

Available MRI events (green) were divided into five modalities (blue) and trimmed incomplete or very poor-quality data (yellow). We further filtered to exclude the relative irrelevant ROIs (orange).
